# Effect of diffusivity of amyloid beta monomers on the formation of senile plaques

**DOI:** 10.1101/2023.07.31.551367

**Authors:** Andrey V. Kuznetsov

## Abstract

Alzheimer’s disease (AD) presents a perplexing question: why does its development span decades, even though individual amyloid beta (Aβ) deposits (senile plaques) can form rapidly in as little as 24 hours, as recent publications suggest? This study investigated whether the formation of senile plaques can be limited by factors other than polymerization kinetics alone. Instead, their formation may be limited by the diffusion-driven supply of Aβ monomers, along with the rate at which the monomers are produced from amyloid precursor protein (APP) and the rate at which Aβ monomers undergo degradation. A mathematical model incorporating the nucleation and autocatalytic process (via the Finke-Watzky model), as well as Aβ monomer diffusion, was proposed. The obtained system of partial differential equations was solved numerically, and a simplified version was investigated analytically. The computational results predicted that it takes approximately 7 years for Aβ aggregates to reach a neurotoxic concentration of 50 μM. Additionally, a sensitivity analysis was performed to examine how the diffusivity of Aβ monomers and their production rate impact the concentration of Aβ aggregates.

## 1. Introduction

Multiple studies have linked the propagation of amyloidogenic proteins to various neurodegenerative conditions like Alzheimer’s disease (AD), Parkinson’s disease, Amyotrophic Lateral Sclerosis, and Huntington’s disease (Jucker & Walker, 2013; Klimova *et al*., 2015). AD is a serious neurodegenerative condition that remains the primary cause of dementia (Hardy, 2006; Maqbool *et al*., 2016; Hung & Fu, 2017). One prominent feature of AD is the spontaneous clustering of amyloid peptides into fibrillar aggregates, driven by a self-sustaining process known as secondary nucleation (Thacker *et al*., 2023). It is widely recognized that individuals with AD display a significant buildup of amyloid beta (Aβ) deposits, also known as senile plaques, in their brain tissue. Yet there remains ongoing debate regarding whether these deposits actively contribute to the onset of the disease or simply signify its advancement (Hardy & Selkoe, 2002; O’Brien & Wong, 2011; Selkoe & Hardy, 2016). Aβ monomers (Aβ40 and Aβ42 peptides) stem from the breakdown of amyloid precursor protein (APP), synthesized in neurons, by β- and γ-secretase enzymes (O’Brien & Wong, 2011; Hampel *et al*., 2021). The majority of Aβ monomers generated are released into the extracellular space, where they have the potential to aggregate (Rahman & Lendel, 2021).

Murphy & Pallitto (2000) investigated the kinetics of Aβ aggregation, while Carbonell *et al*. (2018) provided a comprehensive review of mathematical models addressing protein misfolding mechanisms in neurological diseases. In recent years, the Smoluchowski equations have been used to study Aβ aggregation and diffusion (Achdou *et al*., 2013; Bertsch *et al*., 2017; Bertsch *et al*., 2018; Bertsch *et al*., 2021a; Bertsch *et al*., 2021b; Bertsch *et al*., 2023). The Smoluchowski equations were initially developed in Smoluchowski (1917) to study the rapid coagulation of aerosols. Over time, these equations have been adapted to address various physical scenarios involving the evolving densities of diffusing particles that are prone to coagulation, including protein aggregation (Szała-Mendyk *et al*., 2023). In their general form, the Smoluchowski equations are a set of nonlinear integro-differential equations that do not have a general analytical solution.

An analytical study of the self-assembly processes of filamentous molecular structures, involving primary and secondary nucleation processes and linear growth but excluding the effect of diffusion, was reported in Knowles *et al*. (2009). A minimal model of prion-like brain diseases simulating the production, removal, and transport of toxic proteins in full brain geometry using the Fisher–Kolmogorov-Petrovsky-Piskunov (Fisher–KPP) equation was developed in Weickenmeier *et al*. (2018). The impact of secondary nucleation in Aβ aggregation was examined in Cohen *et al*. (2013). A combined experimental and numerical investigation revealing spatial heterogeneity during protein self-assembly was carried out in Knowles *et al*. (2011). In Morris *et al*. (2008), the Finke-Watzky (F-W) two-step model was employed to fit previously published data on amyloid-beta (Aβ) aggregation, which were reported in Bieschke *et al*. (2005), Vestergaard *et al*. (2005). Typically, data are provided for two isoforms: Aβ40 and Aβ42, with Aβ42 being more prone to aggregation.

The buildup of senile plaques in individuals with AD develops gradually over many years. Interestingly, a study conducted in Meyer-Luehmann *et al*. (2008) revealed that individual plaques can form quite rapidly, within approximately 24 hours. To understand this apparent paradox, this investigation explored the role of Aβ monomer diffusion in the process of Aβ aggregate formation. The proposed hypothesis states that the slow diffusion of Aβ, along with the limited rate of synthesis of Aβ monomers, can be contributing factors to the gradual formation of senile plaques. Thus, it is proposed that the rate of plaque formation is influenced not only by the kinetics of their growth but also by the rate at which the essential reactants (Aβ monomers) are produced and supplied through diffusion, and also by the rate at which Aβ monomers are destroyed by the proteolytic system.

Mathematical models were developed in Kuznetsov & Kuznetsov (2018a), Kuznetsov & Kuznetsov (2018b), Torok *et al*. (2021) to understand the intraneuronal mechanisms driving the formation of amyloid-β plaques. Previous research primarily focused on simulating intracellular processes and neglected the extracellular diffusion of Aβ monomers. Given that Aβ monomers can diffuse very easily, it is important to incorporate this extracellular diffusion process. The current study builds upon recent studies reported in Kuznetsov (2024a), Kuznetsov (2024b), Kuznetsov (2024c), broadening the scope of prior models by focusing on the diffusion of Aβ monomers in the extracellular space and its impact on the formation of Aβ aggregates. The goal is to develop a simple mathematical model for the production, diffusion, and aggregation of Aβ in the context of AD.

## 2. Materials and models

### 2.1. Model equations

The F-W model simplifies the process into two steps, delineating the formation of particular aggregates through two pseudo-elementary reactions: nucleation and autocatalysis. During the nucleation step, new aggregates form continuously, while in the autocatalysis step, these aggregates undergo rapid surface growth (Morris *et al*., 2008; Iashchishyn *et al*., 2017). The two pseudo-elementary reaction steps involve the following:

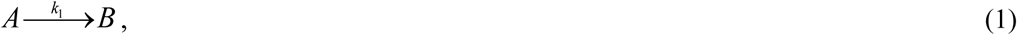

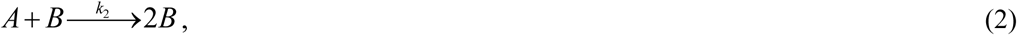

where *A* stands for a monomeric protein, while *B* stands for an amyloid-converted protein. The kinetic constants *k*_1_ and *k*_2_ denote the rates of nucleation and autocatalytic growth, respectively (Morris *et al*., 2008). The primary nucleation process, as described by Eq. (1), involves monomers only. Secondary nucleation, as outlined in Eq. (2), encompasses both monomers and pre-existing aggregates of the same peptide (Thacker *et al*., 2023).

This study employs the F-W model to simulate the conversion of monomers, whose concentration is *C_A_*, into aggregates, whose concentration is represented by *C_B_*. These aggregates include different forms of Aβ oligomers, protofibrils, and fibrils (Chen *et al*., 2017). The buildup of Aβ aggregates results in the formation of amyloid plaques. The simplified nature of the F-W model means it cannot distinguish between different types and sizes of aggregates. Typically, the F-W model is used to study how an initial concentration of monomers transforms into aggregates. In this study, a novel extension of the F-W model is proposed to address scenarios where monomers are continuously introduced through diffusion from the left-hand side of the control volume (CV), represented by the position *x*=0 (Fig. 1). At the boundary *x*=0, which could represent locations such as the cellular membrane (Haass *et al*., 2012; Masters & Selkoe, 2012), Aβ monomers are produced through the cleavage of amyloid precursor protein (APP). The boundary at *x*=*L* represents an extracellular location. The model presented in this paper is given by the following two equations. The first equation is derived by applying the conservation of Aβ monomers within the CV:

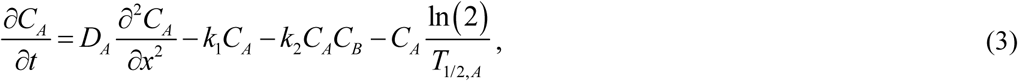

**Fig. 1.**
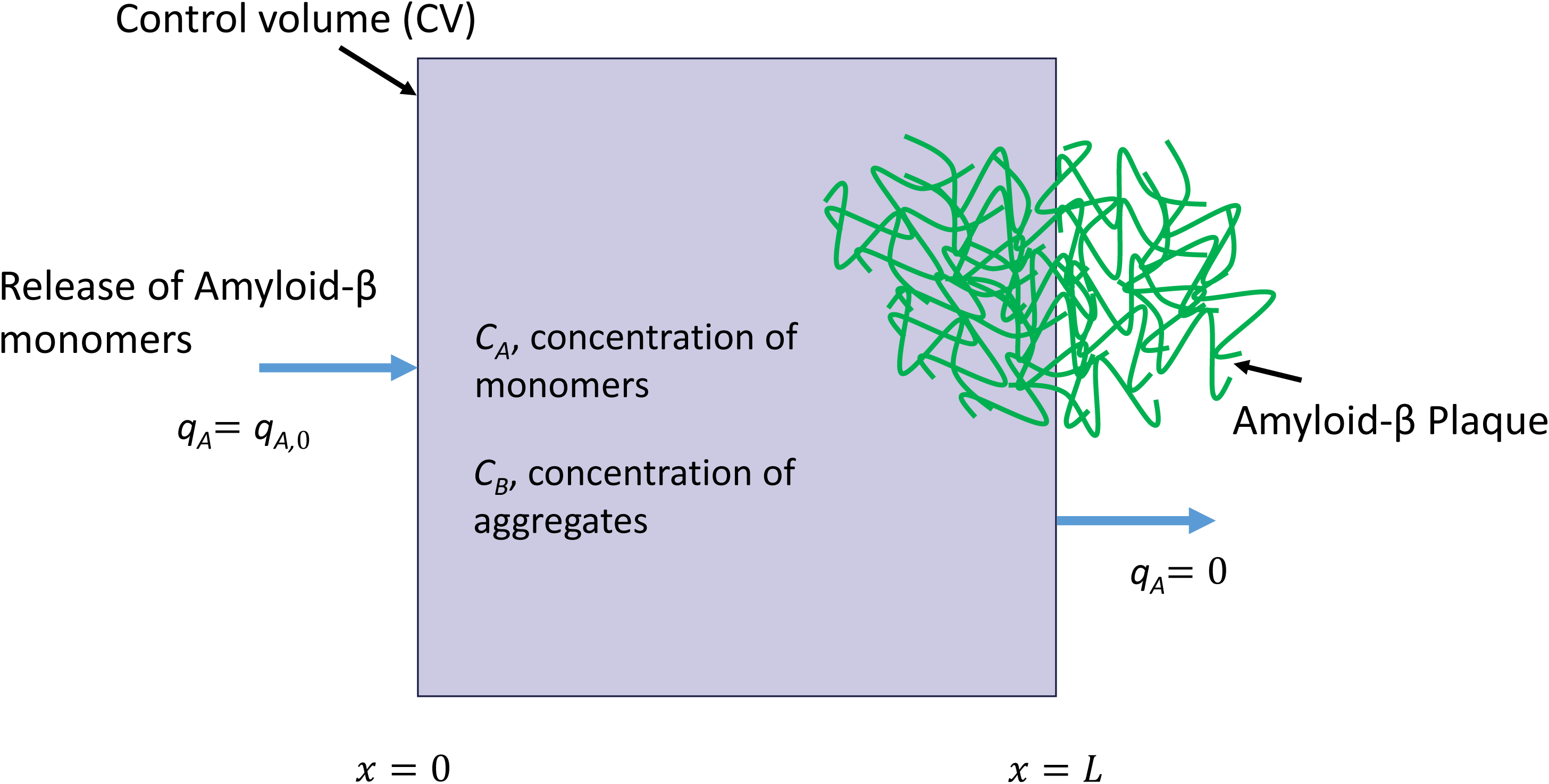
A CV with a width of *L* = 50 μm, used to simulate the diffusion of Aβ monomers from the boundary at *x*=0 and the accumulation of Aβ aggregates. The boundary at *x*=*L* is assumed to be symmetric, indicating that no Aβ monomers pass through it.

where the first term on the right-hand side models the diffusion of Aβ monomers, followed by the second term, which captures the nucleation-driven conversion of these monomers into aggregates. The third term simulates the autocatalytic conversion process of monomers into aggregates, while the fourth term represents the proteolytic degradation of Aβ monomers as described in Saido & Leissring (2012).

The second equation is derived by applying the conservation of Aβ aggregates within the CV:

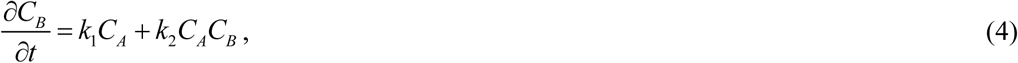

where the first term on the right-hand side represents the nucleation process that converts Aβ monomers into aggregates, while the second term simulates the autocatalytic process responsible for further aggregate formation from monomers.

The initial conditions are outlined below:

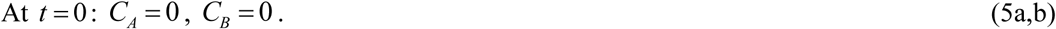

The boundary conditions are outlined below:

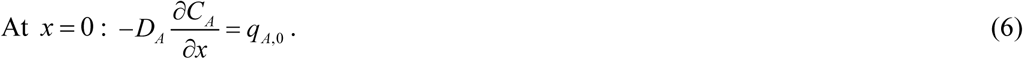

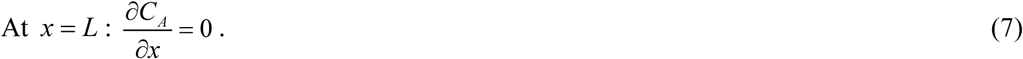

Eqs. (6) and (7) assume the release of Aβ monomers from the boundary situated at *x*=0. The boundary at *x*=*L* is modeled as a symmetric boundary (*L* represents the half distance between senile plaques). Therefore, there is no flux of Aβ monomers through this boundary.

The model utilizes independent variables detailed in Table 1, while Table 2 gives a summary of the dependent variables. Table 3 gives an overview of the parameters used in the model.

**Table 1.** Independent variables utilized in the model.

| Symbol | Definition | Units |
| --- | --- | --- |
| $t$ | Time | s |
| $x$ | Coordinate measuring the distance from the CV's face from which Aβ monomers are supplied | $\mu\text{m}$ |

**Table 2.** Dependent variables utilized in the model.

| Symbol | Definition | Units |
| --- | --- | --- |
| $C_A$ | Concentration of Aβ monomers | $\mu\text{M}$ |
| $C_B$ | Concentration of Aβ aggregates | $\mu\text{M}$ |

**Table 3.**
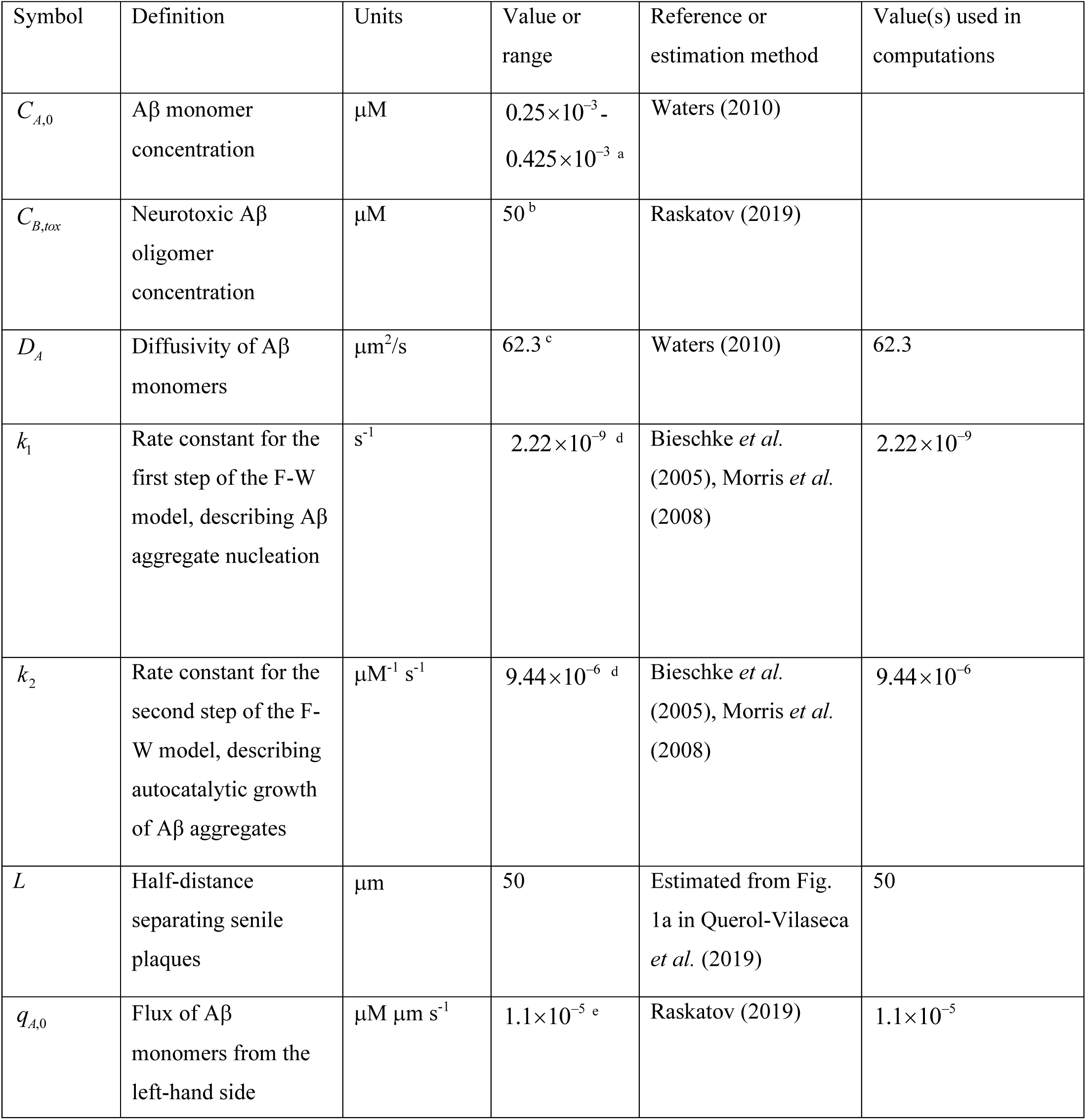
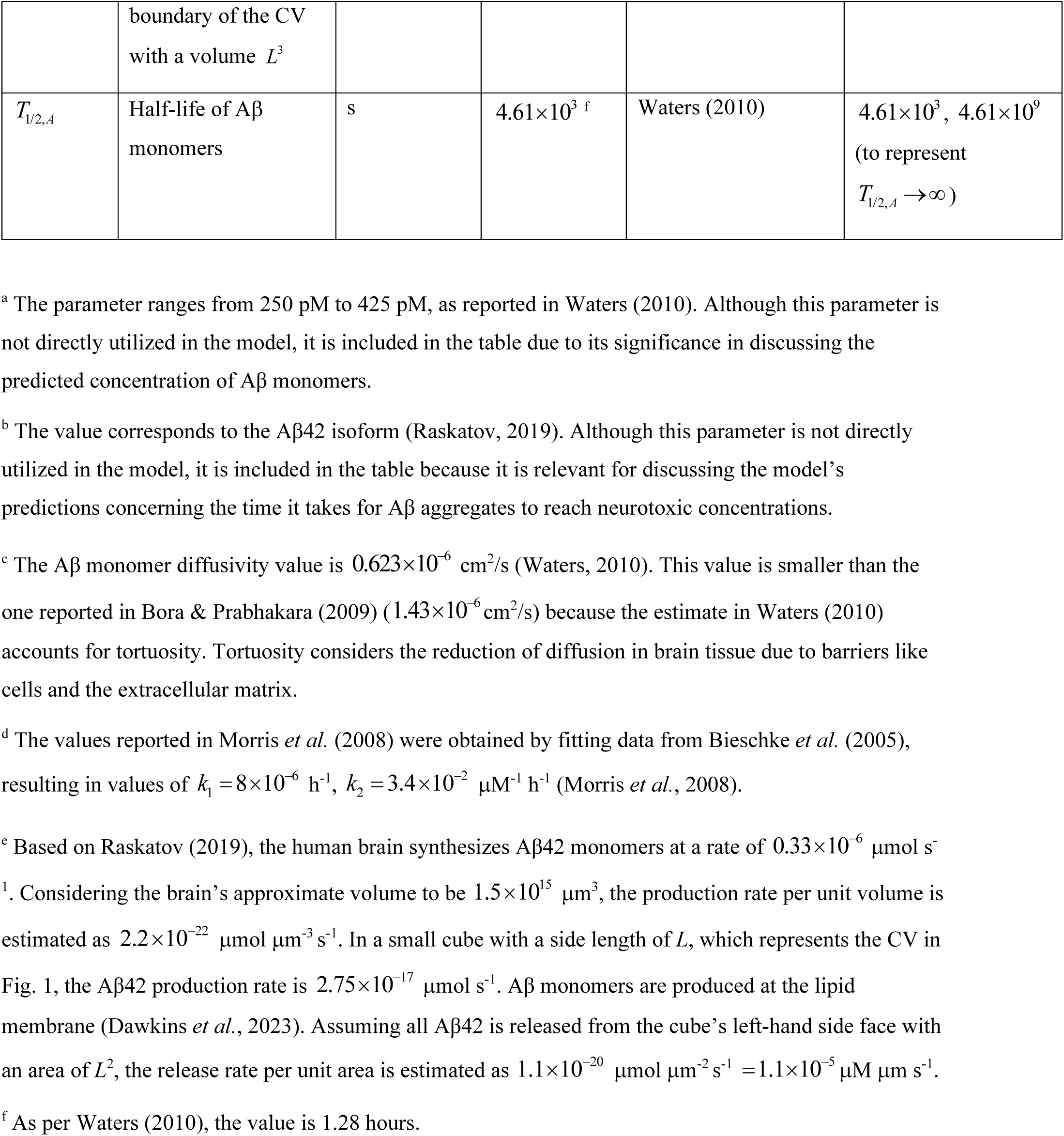
Parameters utilized in the model and the discussion of results.

The dimensionless version of Eqs. (3)-(7) can be found in section S1.1 of the Supplementary Materials. The Aβ aggregation process is affected by three dimensionless parameters: (i) the diffusivity of Aβ monomers, *D*\*_*A*_ = *D*_*A*_ / (*L*^2^*k*_1_); (ii) the production rate of Aβ monomers, *q*\*_*A*,0_ = *q*_*A*,0_*k*_2_ / (*k*^2^_1_*L*) ; and (iii) the half-life of Aβ monomers, *T*\*_1/2,*A*_ =*T*_1/2,*A*_*k*_1_.

If *C_A_* remains nearly constant along *x*, it indicates that the diffusion of Aβ monomers is very fast. One supporting rationale for this assumption is that if soluble Aβ diffuses slower than the rate of aggregation, growth will occur primarily at the tips of the aggregate, leading to a branched, tree-like structure. However, this contradicts experimental observations (Cruz *et al*., 1997), suggesting fast diffusion of Aβ monomers. With this assumption, the CV depicted in Fig. 1 can be regarded as a lumped capacitance body and Eqs. (3) and (4) can be recast as follows:

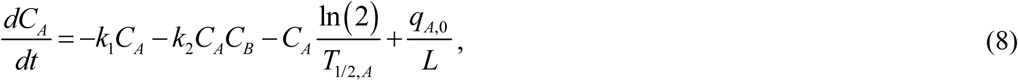

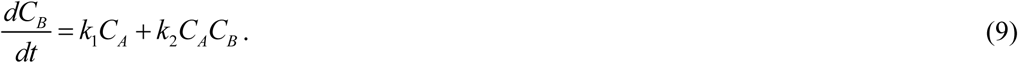

The dimensionless versions of Eqs. (8) and (9) are given in section S1.2 of the Supplementary Materials. When Eqs. (8) and (9) are added, the following result is obtained:

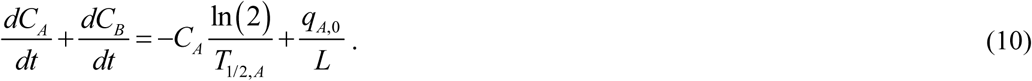

For *T*_1/2,*A*_ →∞, integrating Eq. (10) with respect to time and using the initial condition from Eq. (5a,b) gives the following result:

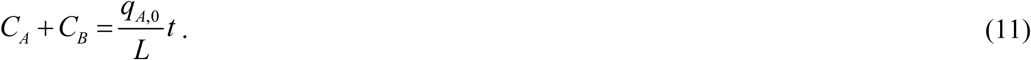

The increase in *C_A_* + *C_B_* over time is due to the flux of Aβ monomers through the left-hand side boundary of the CV. By eliminating *C_A_* from Eq. (4) using Eq. (11), the following equation is obtained:

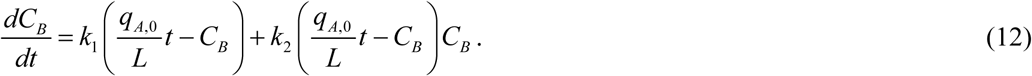

Eq. (12) is similar to the one studied in Kuznetsov & Kuznetsov (2022). To find the exact solution of Eq. (12) with the initial condition given by Eq. (5b), the DSolve function followed by the FullSimplify function in Mathematica 13.3 (Wolfram Research, Champaign, IL) was employed. The resulting solution is as follows:

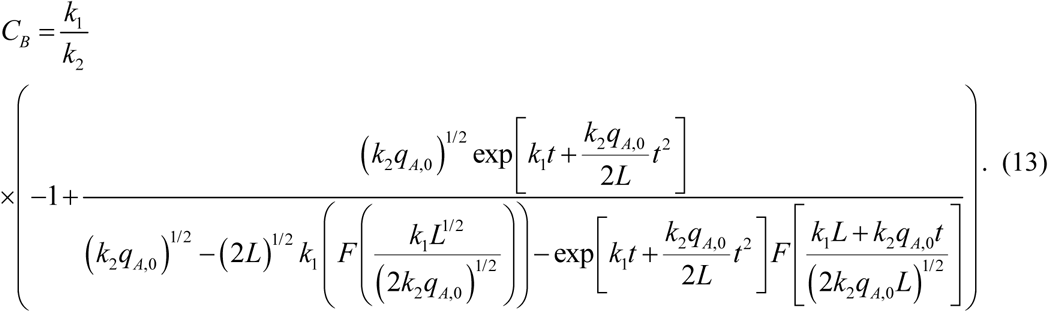

In this equation, *F* (*x*) is Dawson’s integral:

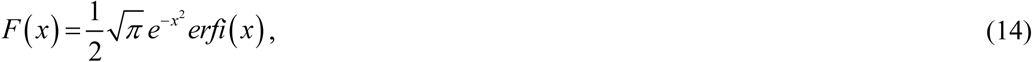

where *erfi* (*x*) is the imaginary error function.

If *t* → 0, Eq. (13) implies that 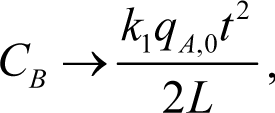, indicating that at small *t*, the concentration of Aβ aggregates is directly proportional to the kinetic constant that describes the nucleation of Aβ aggregates. *C_A_* can then be found using Eq. (11) as:

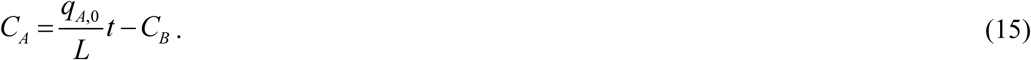

The exact solution provided by Eqs. (13) and (15) is quite cumbersome. A more elegant approximate solution, also valid for the case of *T*_1/2,*A*_ →∞, is

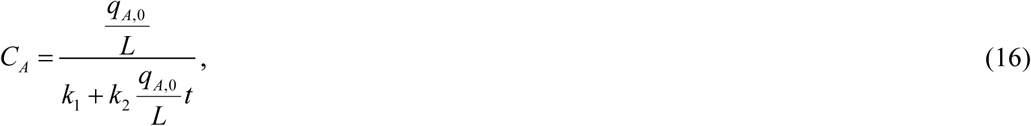

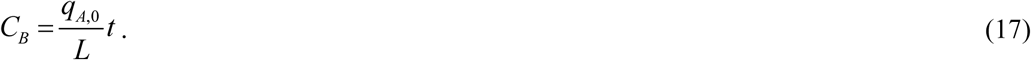

Note that as *t* →∞, Eq. (16) predicts that *C_A_* → 0. The dimensionless forms of Eqs. (16) and (17) are given in section S1.3 in Supplementary Materials.

### 2.2. Sensitivity analysis

One of the primary aims of this research is to examine the sensitivity of Aβ aggregate concentration to variations in the diffusivity of Aβ monomers. The study also explored how changes in the rate of Aβ monomer production, represented by *q_A_*_,0_, affect the concentration of Aβ aggregates.

This was accomplished by calculating the local sensitivity coefficients, which represent the first-order partial derivatives of the observable (the concentration of Aβ aggregates) with respect to the diffusivity of Aβ monomers or the flux of Aβ monomers entering the CV through the left-hand side boundary, following methodologies described in Beck & Arnold (1977), Zadeh & Montas (2010), Zi (2011), Kuznetsov & Kuznetsov (2019). The sensitivity coefficient of *C_B_* to, for instance, *D_A_* was calculated as follows:

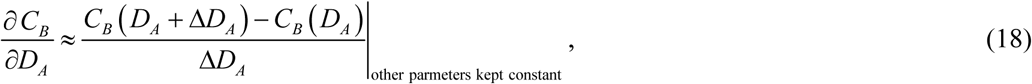

where Δ*D* = 10^−3^ *D_A_* represents the step size. Calculations were conducted using various step sizes to assess the sensitivity coefficients’ independence of the step size.

The non-dimensionalized relative sensitivity coefficient was calculated following the methods outlined in Zadeh & Montas (2010), Kacser *et al*. (1995), as demonstrated below (for example):

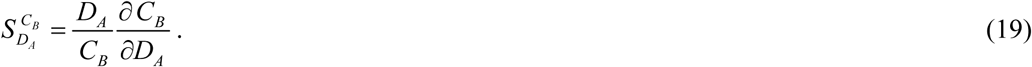

#### S3. Numerical solution

Eqs. (3) and (4) form a system of two partial differential equations (PDEs). They were solved subject to initial conditions given by Eq. (5) and boundary conditions given by Eqs. (6) and (7) using a well-validated MATLAB PDEPE solver (MATLAB R2020b, MathWorks, Natick, MA, USA).

MATLAB’s ODE45 solver was used to solve Eq. (12) with initial condition (5b) (for large values of *t*, using MATLAB’s solver works better than the analytical solution given by Eq. (13)). To ensure accurate results, the error tolerance parameters, RelTol and AbsTol, were set to 1e-10.

## 3. Results

Figs. 2 and 3 depict the scenario where the half-life of Aβ monomers is assumed to be infinite. In Fig. 2a, the computed concentration of Aβ monomers is approximately 0.465×10^−3^ μM, a value nearly within the range reported in Waters (2010) (Table 3): 0.25×10^−3^ to 0.425×10^−3^ μM. The concentrations of monomers, *C_A_*, and aggregates, *C_B_*, gradually decrease from *x*=0 to *x=L*. The concentrations decrease because monomers diffuse from the boundary with the highest concentration (*x*=0) toward the boundary with a symmetric (zero flux) condition (*x*=*L*), where both Aβ monomer and aggregate concentrations reach their lowest levels (see Figs. 2a and 2b). In this scenario, the concentration of aggregates reaches the neurotoxic level of 50 μM (Fig. 2b) within the timescale of *t* = 7.23 years. The model thus recovers an aggregation timescale of 7 years. Interestingly, a simple calculation of the time it takes to form 50 μM of Aβ in the volume under study yields *C_B,tox_L* / *q_A,0_* = 2.27 ×10^8^ s = 7.21 years. This suggests that the timescale for reaching the neurotoxic concentration is primarily governed by the rate of Aβ production.

**Fig. 2.**
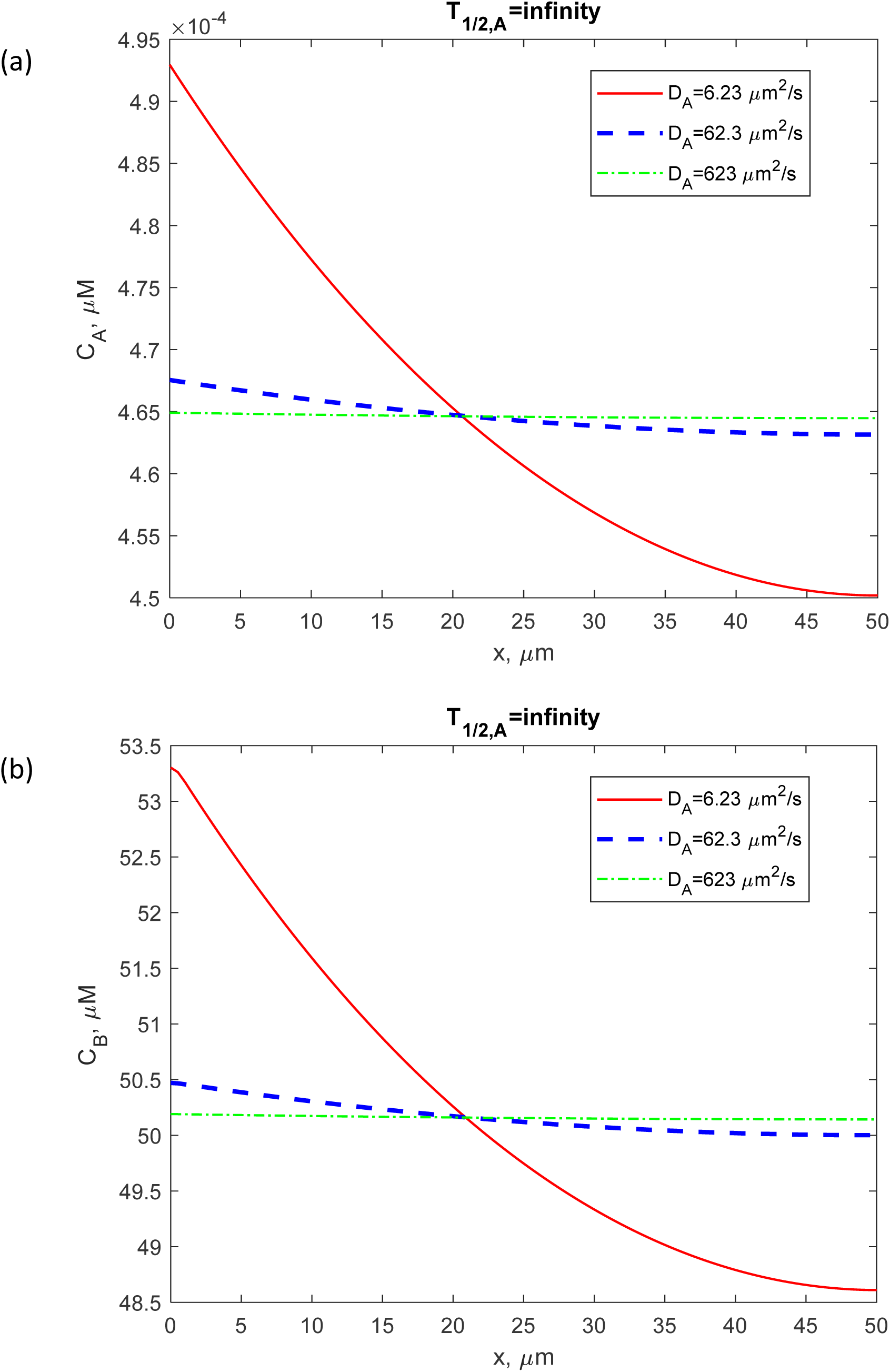
Concentration of Aβ monomers, *C_A_* (a), and concentration of Aβ aggregates, *C_B_* (b), plotted against the distance from the surface releasing Aβ monomers (e.g., cell membrane). These computational results are obtained for *t* = 2.28×10^8^ s = 7.23 years and pertain to the scenario assuming an infinite half-life of Aβ monomers.

**Fig. 3.**
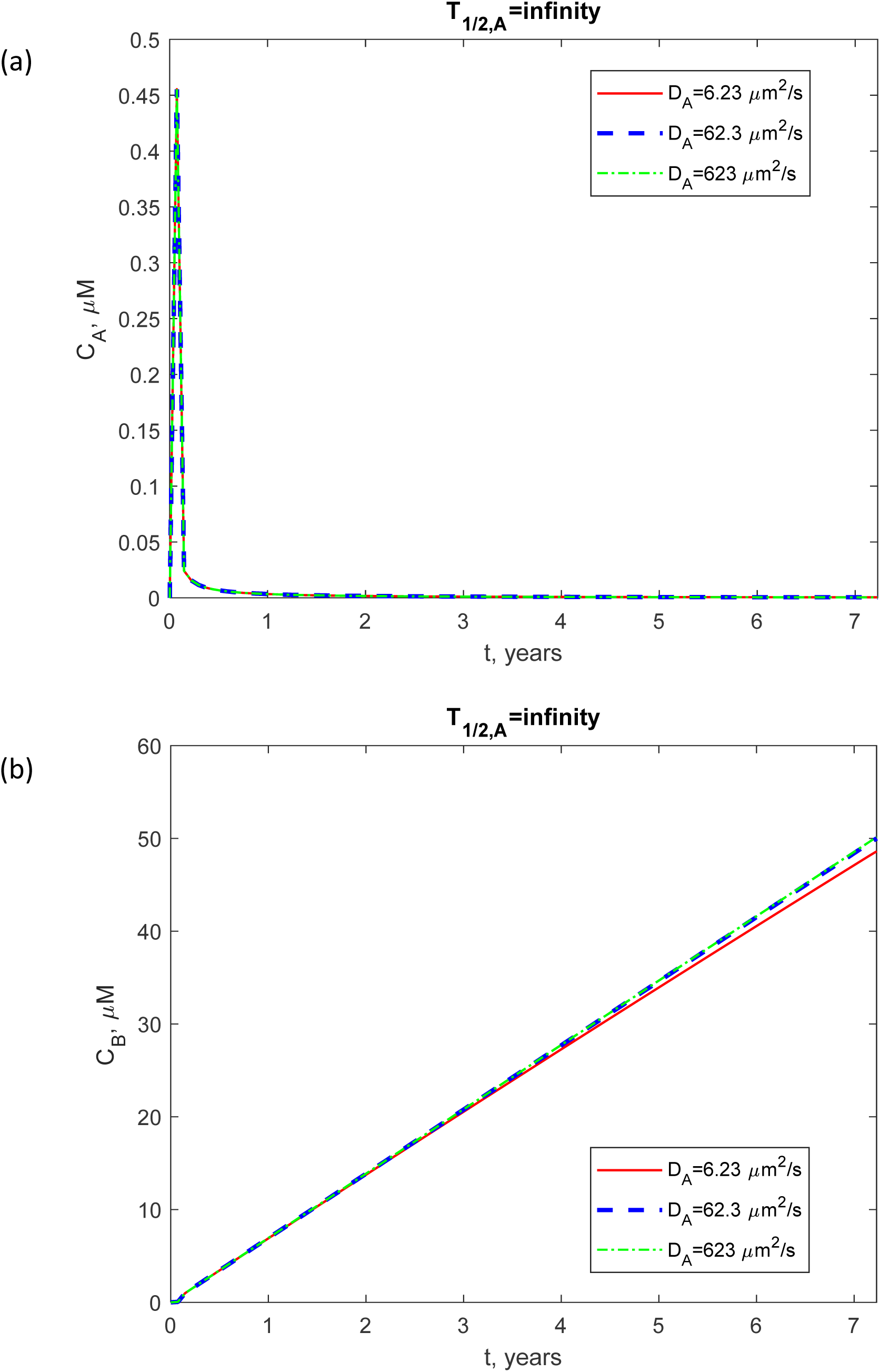
(a) Concentration of Aβ monomers, *C_A_*, plotted against time (in years). (b) The concentration of Aβ aggregates, *C_B_*, also plotted against time (in years). The computational results are presented at the right-hand side boundary of the CV, *x* = *L*. This scenario assumes an infinite half-life of Aβ monomers.

Initially, at the right-hand side boundary of the CV, *x* = *L*, the concentration of Aβ monomers is zero, as indicated by the initial condition in Eq. (5a). Subsequently, it rapidly increases, followed by a decline, and then stabilizes at a low level due to the conversion of monomers into aggregates (Fig. 3a). The concentration of aggregates steadily increases in a linear fashion owing to the continuous supply of reactants (monomers) through the left-hand side boundary of the CV at *x*=0 (Fig. 3b). When the diffusivity of the monomers is decreased, the concentration of aggregates exhibits a slightly slower increase, as illustrated by the red solid curve in Fig. 3b corresponding to *D_A_* = 6.23 μm²/s.

To gain a deeper understanding of how the concentrations of Aβ monomers and aggregates evolve over time, a simplified lumped capacitance model given by Eqs. (8) and (9) was developed. In Fig. S1, presented in the Supplementary Materials, the numerical solution of Eqs. (8) and (9) for the case of *T*_1/ 2, *A*_ →∞ was compared to the approximate analytical solution given by Eqs. (16) and (17). Remarkably, the approximate analytical solution of the lumped capacitance model shows excellent agreement with the numerical solution. As time progresses, the concentration of Aβ monomers decreases proportionally to 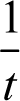 (Eq. (16)). This is consistent with the F-W model, which disregards any potential equilibrium state between the monomers and aggregates, and predicts that the aggregation continues until all monomers are converted into aggregates (Martin, 2020; Finke *et al*., 2020). The concentration of Aβ aggregates increases linearly with time (Eq. (17)). This is due to the reactants (Aβ monomers) entering the CV (Fig. 1) at a constant rate of *q_A,0_L*^2^; all reactants are subsequently converted into Aβ aggregates.

For a finite half-life of Aβ monomers, *T*_1/2,*A*_ = 4.61×10^3^ s (Figs. 4 and 5), the concentration of Aβ aggregates is reduced significantly compared to the scenario with an infinite half-life, decreasing from approximately 50 μM to around 5.25×10^−3^ μM (compare Fig. 4b and Fig. 2b). The significant drop in aggregate concentration occurs because, with a shorter half-life, a considerable portion of the monomers are broken down by the proteolytic system instead of forming aggregates. This finding has important implications for AD. Numerous studies have associated AD with the malfunctioning of the proteolytic machinery, leading to the inability to regulate Aβ concentrations effectively and degrade misfolded Aβ, which has a propensity to aggregate (Wang *et al*., 2006; Baranello *et al*., 2015; Kim & Priefer, 2020). The study’s findings suggest that Aβ aggregation can still occur even when the Aβ monomer concentration is relatively low, provided that the protein clearance mechanism fails to continuously degrade newly generated Aβ monomers. Preventing Aβ aggregation requires more than just breaking down faulty Aβ. Conversely, a well-functioning proteolytic system capable of degrading Aβ monomers can protect the brain from Aβ aggregates. Interestingly, achieving this protection does not require the ability to degrade aggregates (notably, Eq. (4) lacks a degradation term).

**Fig. 4.**
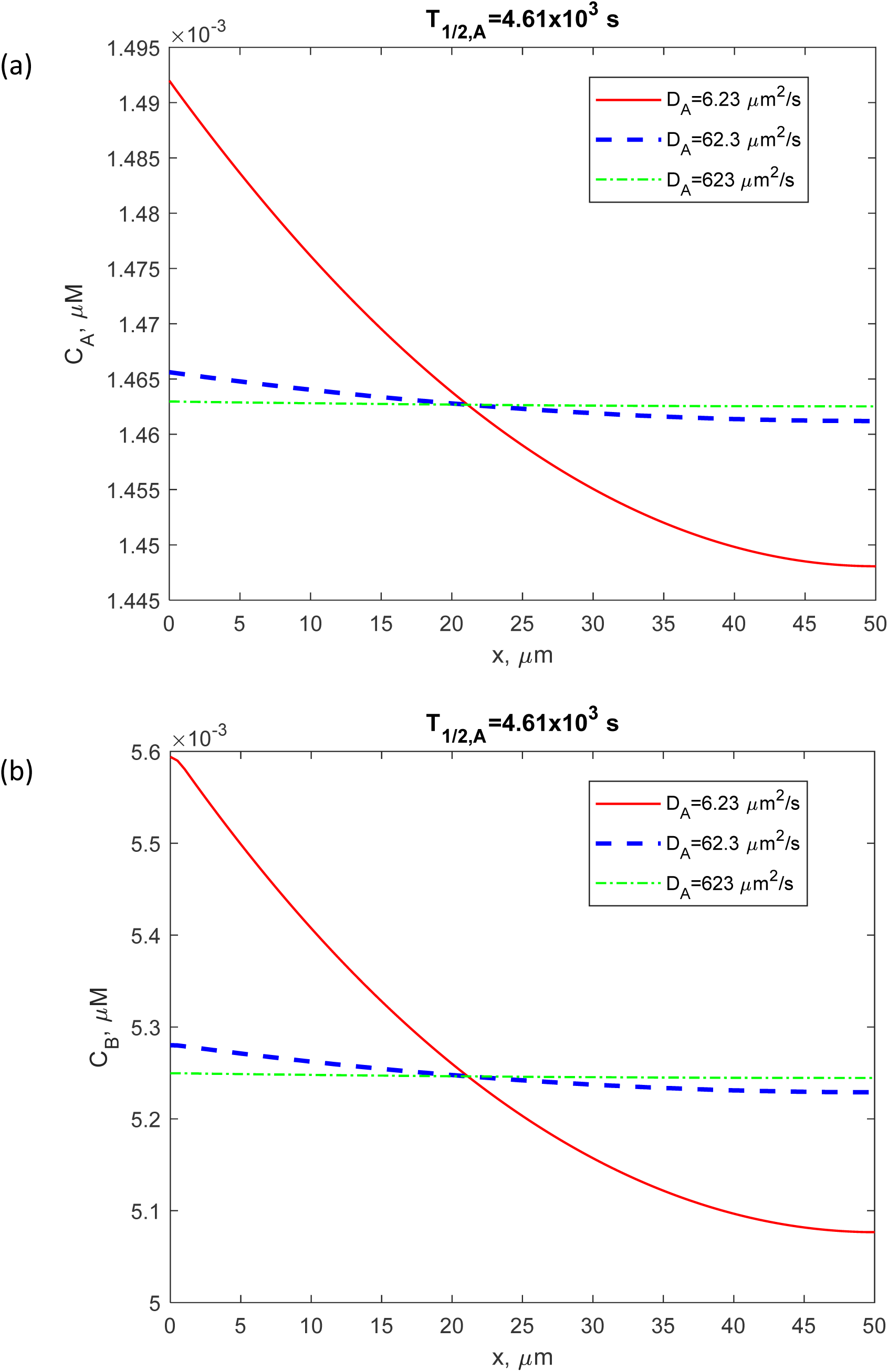
(a) The concentration of Aβ monomers, *C_A_*, as a function of the distance from the surface releasing Aβ monomers (such as the cell membrane). (b) The concentration of Aβ aggregates, *C_B_*, as a function of the distance from the same surface. The computational results are displayed at *t* = 2.28×10^8^ s = 7.23 years. This figure illustrates a scenario where Aβ monomers have a finite half-life (*T* = 4.61×10^3^ s).

**Fig. 5.**
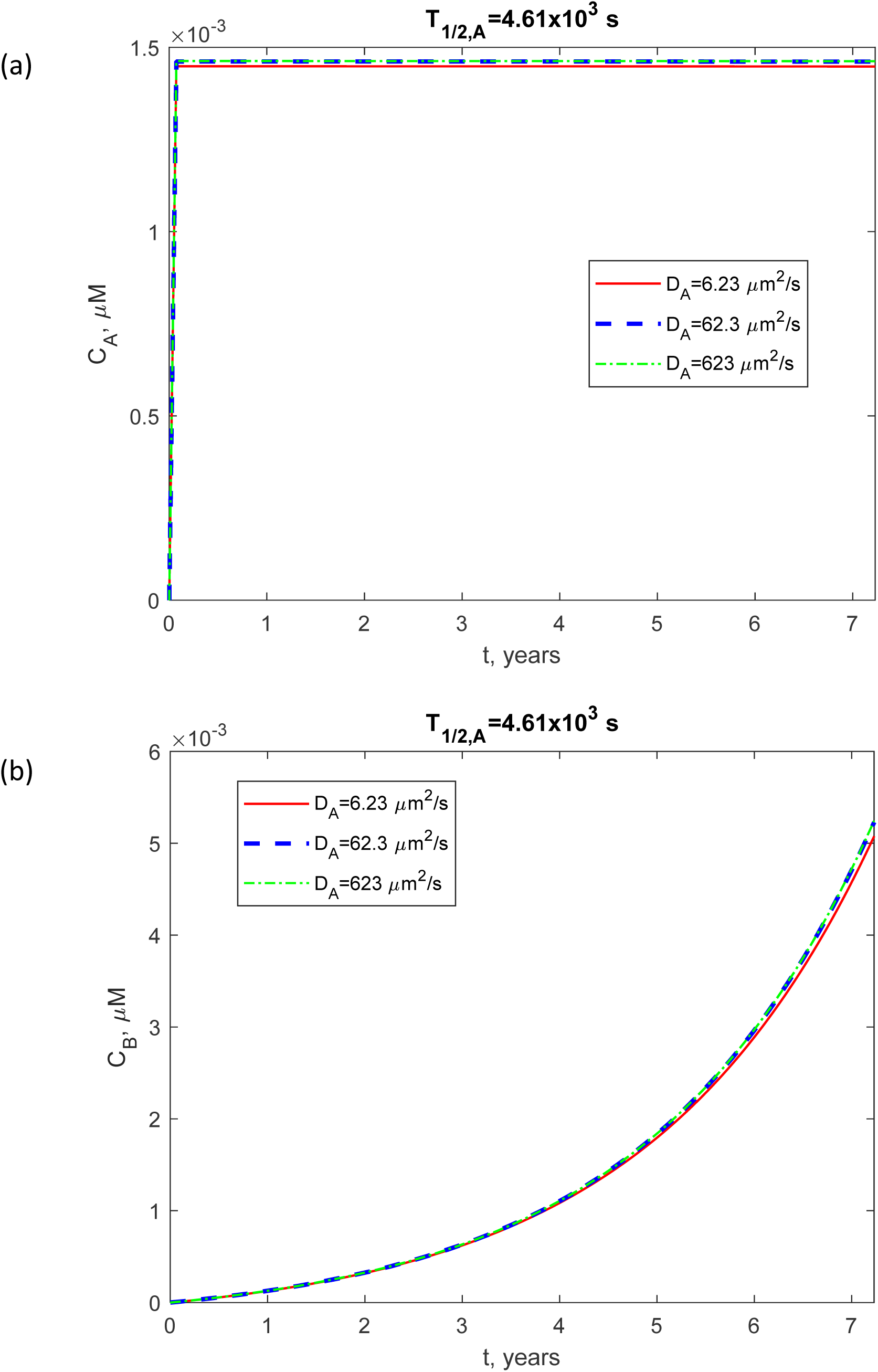
(a) Concentration of Aβ monomers, *C_A_*, plotted against time (in years). (b) Similarly, the concentration of Aβ aggregates, *C_B_*, depicted over time. The computational results are depicted at the right-hand side boundary of the CV, *x* = *L*. This scenario assumes a finite half-life of Aβ monomers (*T* = 4.61×10^3^ s).

When simulating the scenario with a finite half-life of Aβ monomers, which corresponds to a well-functioning Aβ monomer clearance system, the initial peak in the concentration of Aβ monomers is absent (compare Fig. 5a and Fig. 3a). Furthermore, the increase in the concentration of Aβ aggregates, *C_B_*, does not follow a linear trend and occurs much slower, ultimately reaching a significantly lower value after 7 years (compare Fig. 5b and Fig. 3b). Since the concentration of Aβ aggregates in Fig. 5b is four orders of magnitude lower than the neurotoxic level, while it reaches the neurotoxic level in Fig. 3b, the malfunctioning of the machinery responsible for degrading Aβ monomers may be a necessary condition for developing AD.

When the diffusivity of Aβ monomers, *D_A_*, is increased, the concentration of Aβ monomers, *C_A_*, initially exhibits an increase. However, once *D_A_* exceeds 30 μm²/s, *C_A_* reaches a plateau and remains constant even with further increases in *D_A_* (Fig. 6a). A similar trend is observed for the concentration of Aβ aggregates, *C_B_* (Fig. 6b). This suggests that when the diffusivity of Aβ monomers exceeds 30 μm²/s, the diffusion of Aβ monomers no longer presents a limiting factor for Aβ aggregation.

**Fig. 6.**
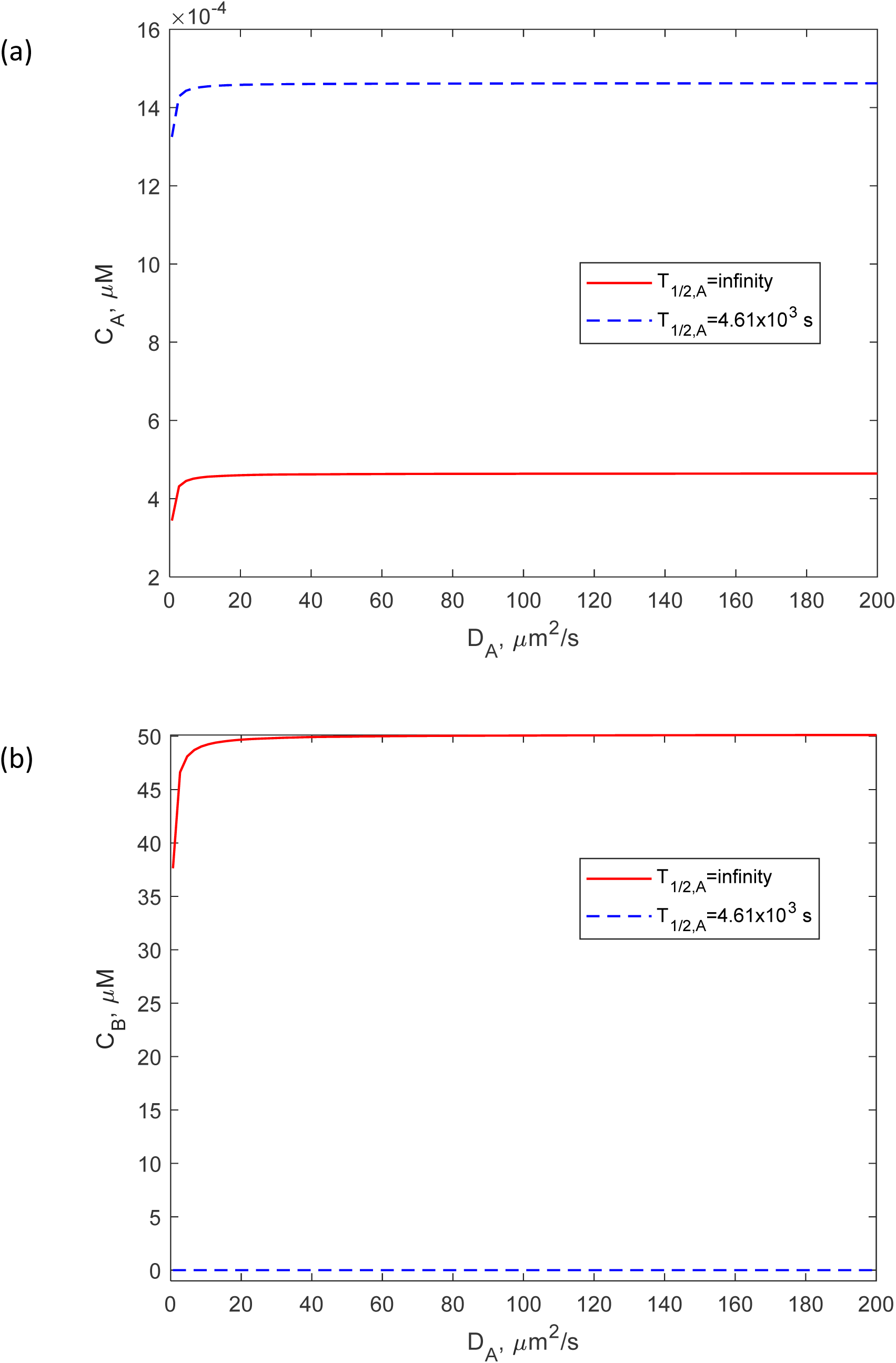
(a) Concentration of Aβ monomers, *C_A_*, plotted against the diffusivity of Aβ monomers, *D_A_*. (b) Concentration of Aβ aggregates, *C_B_*, also plotted against the diffusivity of Aβ monomers, *D_A_*. The computational results are shown at the right-hand side boundary of the CV, *x=L*, and *t* = 2.28×10^8^ s = 7.23 years.

Interestingly, in Fig. 6a, the concentration of Aβ monomers, *C_A_*, for the scenario with an infinitely large half-life, is lower than the concentration for the case with a finite half-life of Aβ monomers. It is important to note that the distributions of *C_A_* shown in Fig. 6a are not steady-state distributions of monomer concentration; the steady-state monomer concentration is zero (as indicated by Eq. (16)). Fig. 6a is computed for *t* = 7.23 years, and if time is increased further, the monomer concentrations for both infinite and finite half-life scenarios will eventually decay to zero. In Fig. 6b, it is evident that the concentration of aggregates is larger for the scenario with an infinite half-life of monomers, which is consistent with expectations.

It is intriguing that in the scenario with an infinite half-life of Aβ monomers, the concentration of Aβ monomers, *C_A_*, appears to be nearly independent of the rate of Aβ monomer production, *q_A_*_,0_, as indicated by the horizontal solid red line in Fig. 7a. To explore this further, a model obtained under the lumped capacitance approximation was employed. This model (see Eqs. (8) and (9)) assumes a lack of concentration variations, *C_A_* and *C_B_*, with respect to *x*; it accounts for concentrations changing with time but not with the distance from the *x*=0 plane. Fig. S1 explores a hypothetical scenario covering 70 years. Based on Fig. 6, the threshold value (30 μm^2^/s) is smaller than the value used in the calculations (*D_A_* = 62.3 μm^2^/s); therefore, the spatial heterogeneity of Aβ distribution can be neglected, and the use of the lumped capacitance approximation is not expected to affect the accuracy of the results. Fig. S1a shows that *C_A_* becomes small after a short transient and decreases to zero as time progresses. Since the lumped capacitance model assumes an infinitely large value of Aβ monomer diffusivity, this suggests that *C_A_* is primarily determined by the difference between the production rate of Aβ monomers and their conversion rate into aggregates. The concentration of Aβ aggregates, *C_B_*, exhibits a linear increase with time (Fig. S1b). This is attributed to a constant rate of Aβ monomer production, *q_A_*_,0_, which enables continuous conversion of Aβ monomers into aggregates. Similar trends can be observed in Fig. 3b, where the finite rate of diffusion of Aβ monomers is taken into account.

**Fig. 7.**
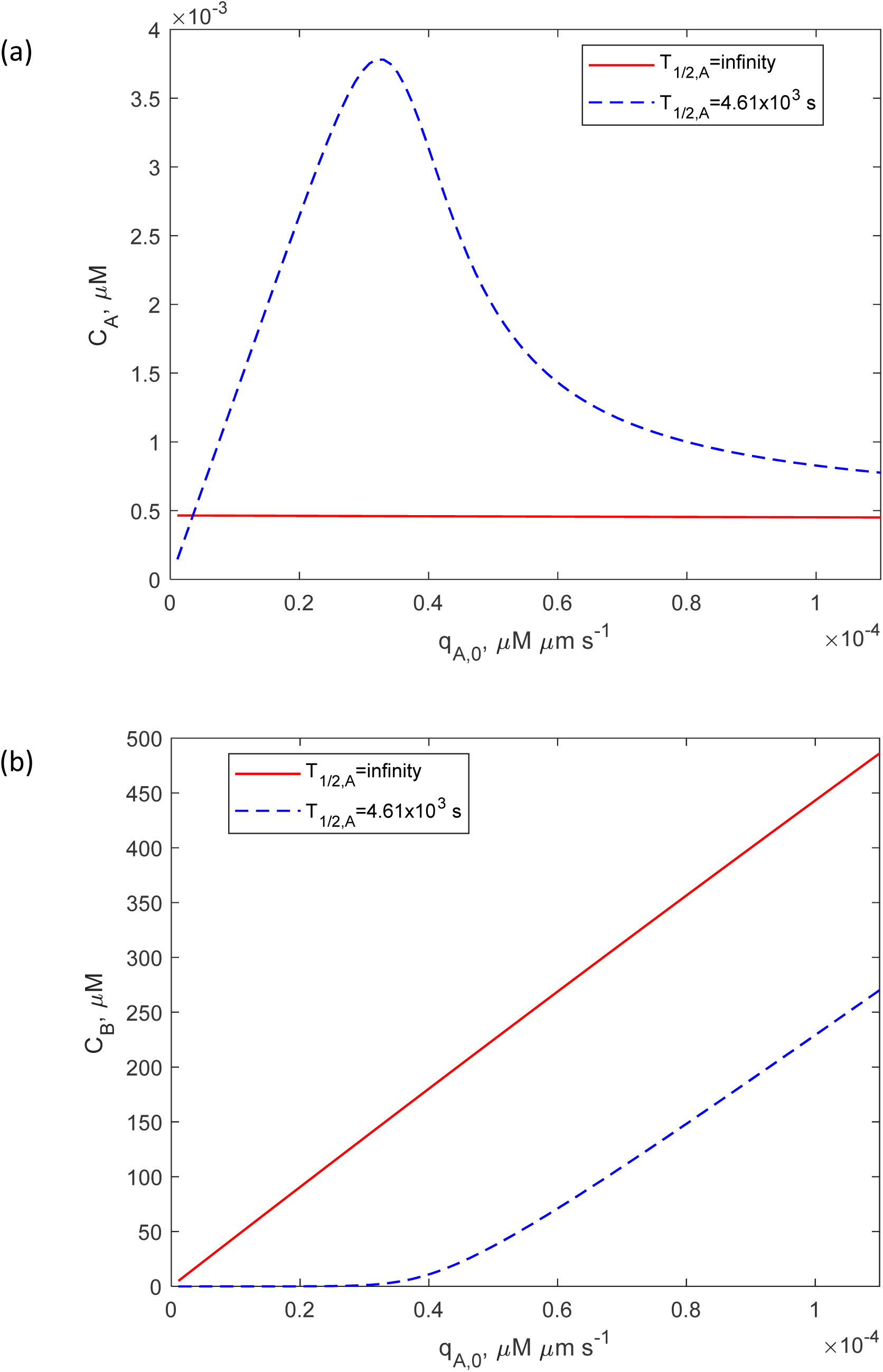
(a) The concentration of Aβ monomers, *C_A_*, plotted against the flux of Aβ monomers from the left-hand side boundary of the CV, *q_A_*_,0_. (b) The concentration of Aβ aggregates, *C_B_*, also plotted against the flux of Aβ monomers from the right-hand side boundary of the CV, *q_A_*_,0_. The computational results are shown at the boundary *x=L* and *t* = 2.28×10^8^ s = 7.23 years.

As depicted in Fig. 7b, for an infinite half-life of Aβ monomers the concentration of Aβ aggregates, *C_B_*, at the boundary *x=L* (50 μm) and *t* = 7.23 years, increases linearly with *q_A_*_,0_ at large times. This behavior is attributed to the fact that *q_A_*_,0_ represents the rate at which reactants are produced and subsequently converted into Aβ aggregates. The production rate, *q_A_*_,0_, is assumed to be constant and independent of time. When the half-life of Aβ monomers is finite (4.61×10^3^ s), an increase in *q* results in a delayed increase in *C_B_*, as long as *q_A_*_,0_ remains small. However, once *q_A_*_,0_ exceeds 0.4 μMꞏμmꞏs⁻¹, *C_B_* begins to increase at the same rate as it does in the case with an infinite half-life of Aβ monomers (Fig. 7b).

The dimensionless sensitivity of the molar concentration of Aβ aggregates, *C_B_*, to the diffusivity of Aβ monomers, *D_A_*, remains consistently positive at all times *t* (Fig. 8a). This is because Aβ monomers serve as reactants in the F-W model, which describes Aβ aggregation (Eqs. (1) and (2)). An increase in the diffusivity of monomers enhances the supply of reactants (see Figs. S2 and S3 in the Supplementary Materials, showing the flux of Aβ monomers), resulting in a greater accumulation of aggregated Aβ.

**Fig. 8.**
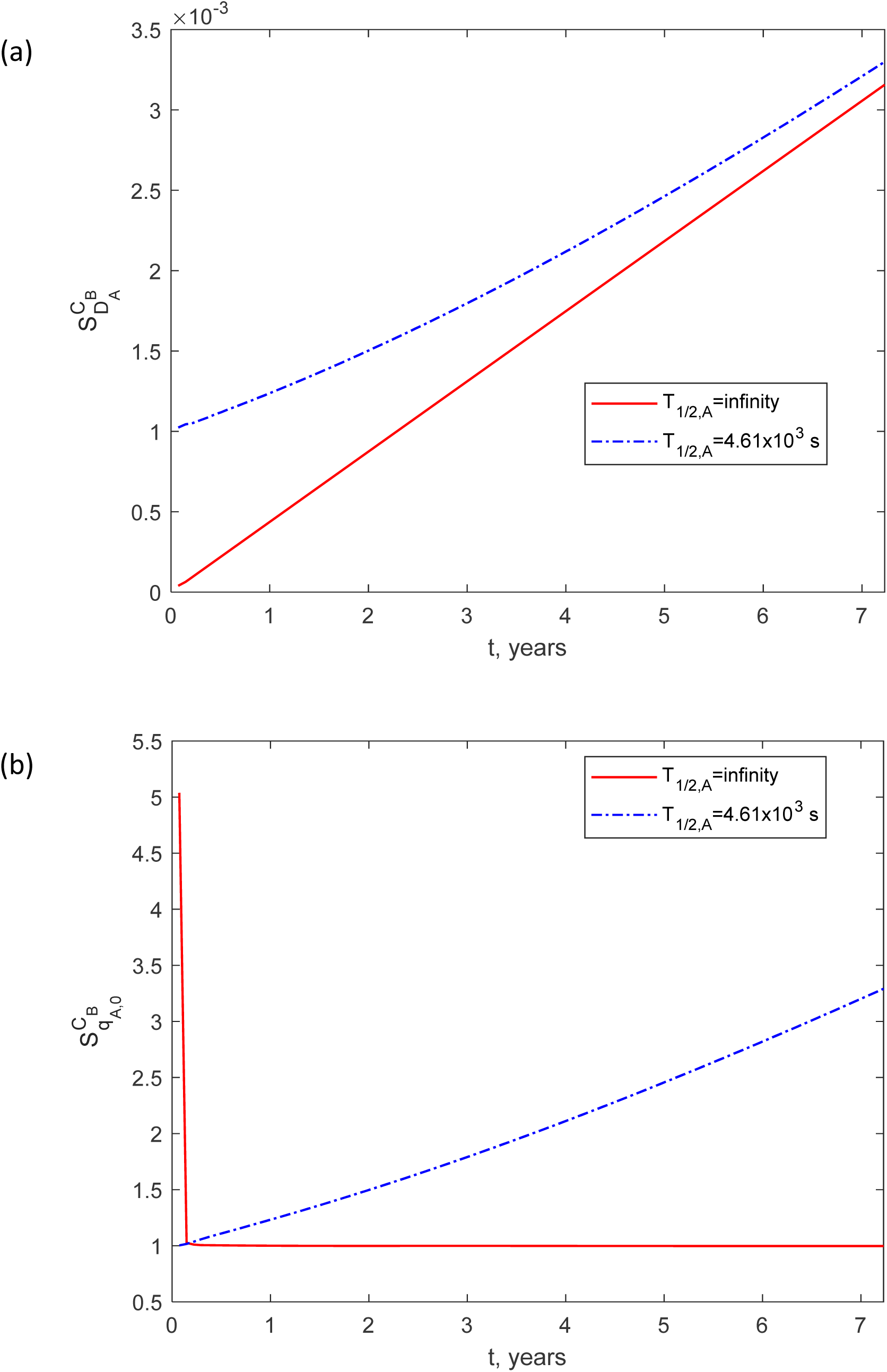
(a) The dimensionless sensitivity of the molar concentration of Aβ aggregates, *C_B_*, to the diffusivity of Aβ monomers, *D_A_*, over time. (b) The dimensionless sensitivity of *C_B_* to the flux of Aβ monomers from the left-hand side boundary of the CV, *q_A_*_,0_, over time. The calculations were carried out for Δ*D_A_* = 10^−3^ *D_A_*, and close results were obtained for Δ*D_A_* = 10^−2^ *D_A_*. The computational findings are displayed at the right-hand side boundary of the CV, *x* = *L*.

The dimensionless sensitivity of the concentration of Aβ aggregates, *C_B_*, to the rate of Aβ monomer production, *q_A_*_,0_, which enters the model through the flux of Aβ monomers from the left-hand side boundary, is also positive at all times *t* (Fig. 8b). It is worth noting that in the scenario with an infinite half-life of Aβ monomers (represented by the red line in Fig. 8b), the sensitivity of *C_B_* to *q_A_*_,0_ quickly stabilizes at a constant value of 1. This behavior is attributed to the linear dependence of *C_B_* on *q_A_*_,0_ (as evident from the red solid line in Fig. 7b), indicating that *C_B_* is directly influenced by the rate of production of the reactants (Aβ monomers). The value of 1 in the sensitivity calculation is a result of rescaling in Eq. (19) that is performed to ensure that the dimensionless sensitivity is independent of magnitudes of parameters.

## 4. Discussion, limitations of the model, and future directions

The findings of this paper indicate that the diffusion of Aβ monomers becomes a limiting factor when their diffusivity is below 30 μm^2^/s. Therefore, for physiologically relevant diffusivity values of Aβ monomers, the formation of Aβ aggregates is primarily governed by the production and removal rates of Aβ.

A popular hypothesis suggests that sporadic forms of AD are linked to impaired clearance of Aβ aggregates (Selkoe, 2001; Tanzi *et al*., 2004; Saido & Leissring, 2012). A modeling paper that utilized the Smoluchowski equations found that an increased removal rate of toxic Aβ results in a broader healthy region of the brain (Bertsch *et al*., 2021a). The findings of this paper suggest that preventing AD may not hinge solely on the proteolytic system’s capability to break down Aβ polymers. The results indicate that even if the proteolytic system only breaks down Aβ monomers, the aggregation of Aβ cannot advance due to the insufficient availability of reactants (monomers). This insight has significant implications for potential therapeutic approaches to AD.

The current model has limitations, notably because the F-W model it utilizes does not account for the size variability of Aβ polymers. It simplifies the Aβ size distribution to just monomers and aggregates. Future work should incorporate models, like those based on the Smoluchowski equations, which describe the concentrations of aggregates of varying sizes as they combine to form larger clusters. However, it is unlikely that incorporating the size distribution of aggregates would alter the main results.

Future studies should also focus on developing models that accurately depict the interaction between Aβ aggregates and tau tangles (Hampel *et al*., 2021). To tackle this issue, the model should incorporate cross-membrane diffusion processes, considering that Aβ aggregates primarily form in the extracellular space, while tau tangles are mainly intracellular. Moreover, future modeling should consider the activation of microglia in response to Aβ aggregation. Exploring the potential synergy between microglia-mediated degradation of Aβ aggregates and the neuroinflammatory response induced by microglia activation (Hansen *et al*., 2018; Chen *et al*., 2017; Meyer-Luehmann *et al*., 2008) is crucial.

Future extensions of the model should consider that Aβ transport in the extracellular space may be influenced not only by diffusion but also by convection and other active transport mechanisms. While it could be argued that Aβ, potentially being a waste product, might evade active transport processes and that its spatial distribution is primarily affected by diffusion, the use of Aβ levels and Aβ40/Aβ42 ratios in blood as important clinical markers for AD suggests that peripheral Aβ concentrations reflect those in the brain.

To improve the accuracy of the proposed model’s predictions, future experimental results are needed to better estimate the values of the parameters involved in the model. Additionally, further research should focus on extending the developed model to include the interaction between Aβ peptides and tau proteins, following the path suggested in the studies by Bertsch *et al*. (2021a), Bertsch *et al*. (2021b), Bertsch *et al*. (2023).

### Ethical statement

The manuscript does not contain human or animal studies.

## Abbreviations

Aβ: amyloid beta
AD: Alzheimer’s disease
CV: control volume
F-W: Finke-Watzky

## Supplemental Materials

### S1. Dimensionless equations

#### S1.1. Dimensionless form of **Eqs. (3)**-(7)

Eqs. (1)-(7) are converted into their dimensionless form as follows:

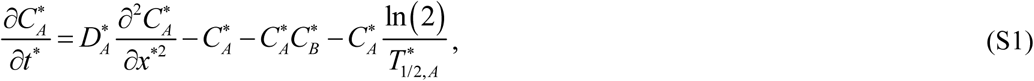

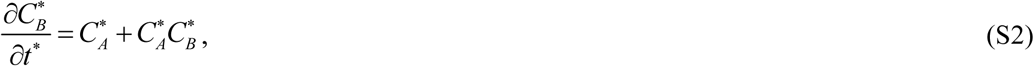

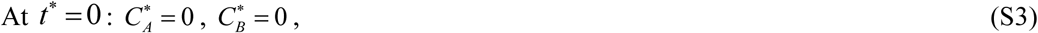

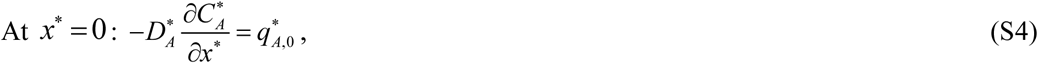

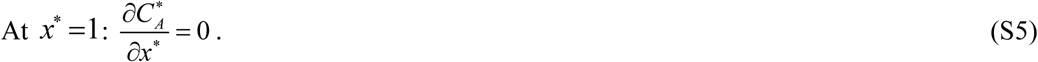

#### S1.2. Dimensionless form of Eqs. (8) and (9)

The dimensionless form of Eqs. (8) and (9) is as follows:

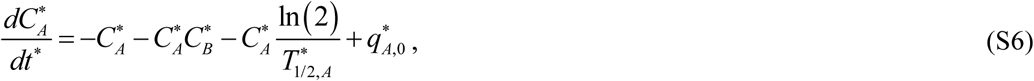

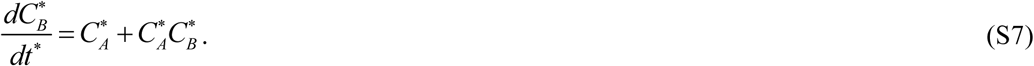

Eqs. (S6) and (S7) should be solved with the initial conditions provided in Eq. (S3).

#### S1.3. Dimensionless form of Eqs. (16) and (17)

The dimensionless form of Eqs. (16) and (17) is as follows:

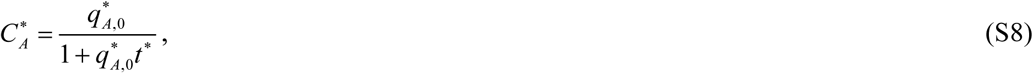

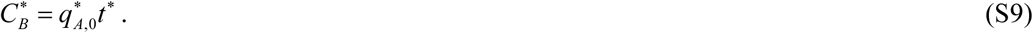

### S2. Tables

**Table S1.** Dimensionless independent variables used in the model.

| Symbol | Definition |
| --- | --- |
| $t^*$ | $t k_1$ |
| $x^*$ | $x / L$ |

**Table S2.** Dimensionless dependent variables used in the model.

| Symbol | Definition |
| --- | --- |
| $C_A^*$ | $C_A k_2 / k_1$ |
| $C_B^*$ | $C_B k_2 / k_1$ |

**Table S3.**
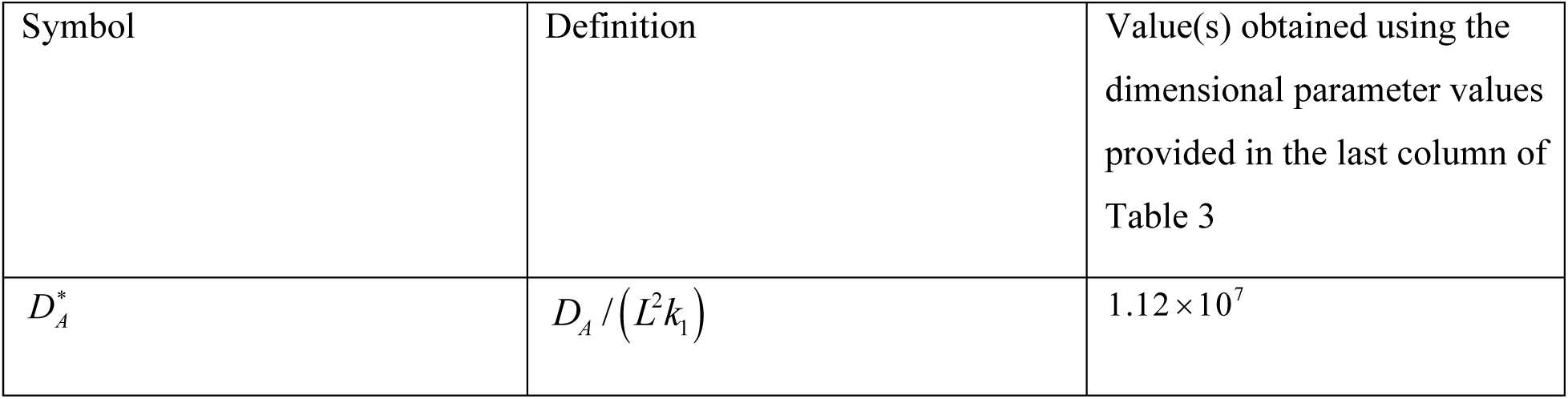
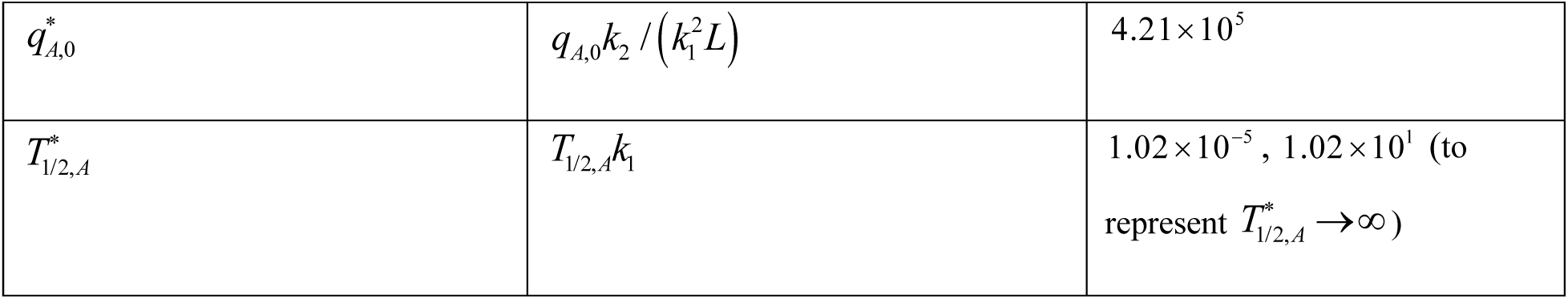
Dimensionless parameters used in the model.

### S3. Supplementary figures

**Fig. S1.**
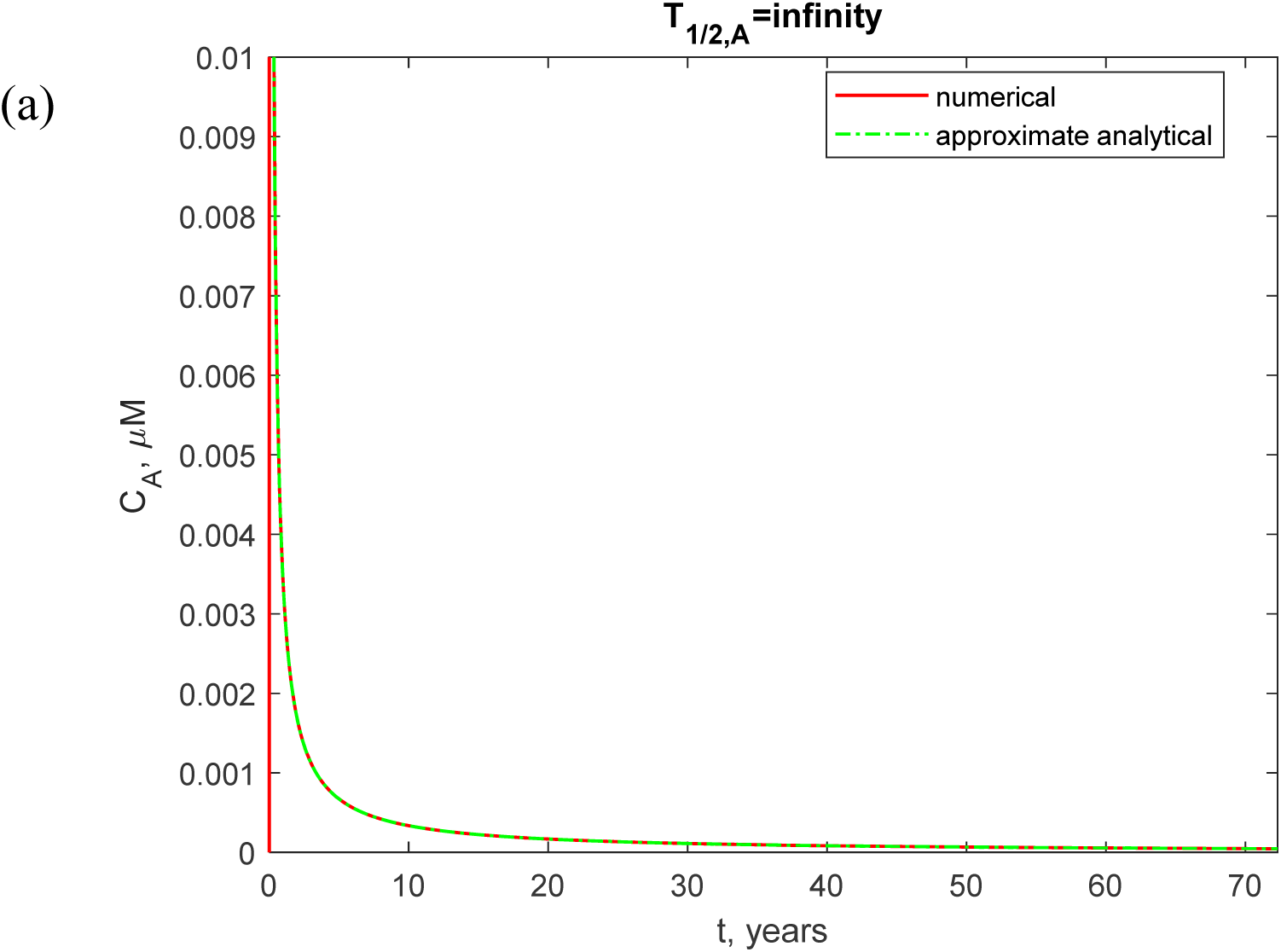
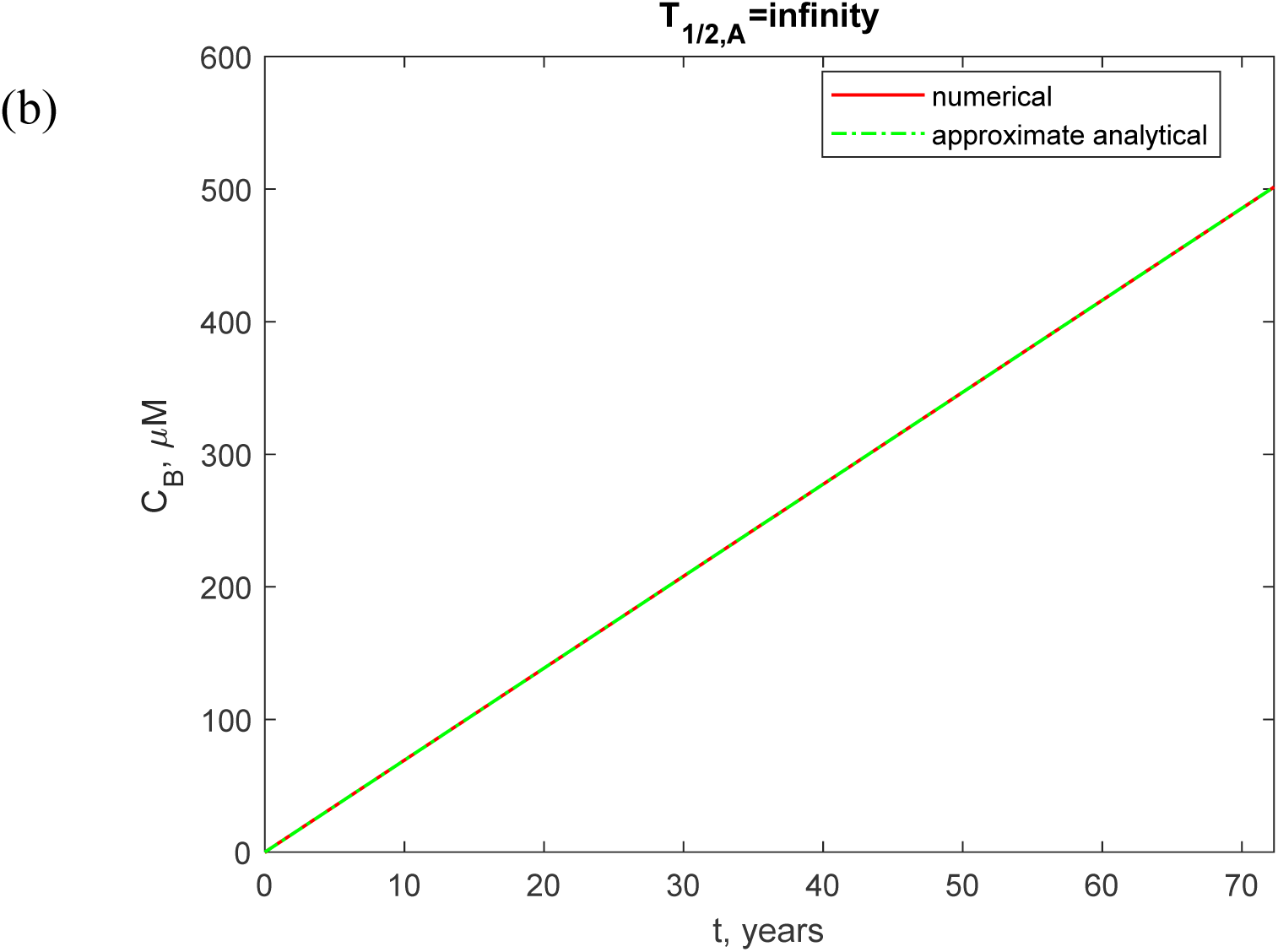
A comparison between the numerical solutions of Eqs. (8) and (9) and the approximate solutions given by Eqs. (16) and (17). (a) Molar concentration of Aβ monomers, *C_A_*, as a function of time (in years). (b) Molar concentration of Aβ aggregates, *C_B_*, as a function of time (in years). A hypothetical scenario spanning 70 years. *D_A_* = 62.3 μm^2^/s.

**Fig. S2.**
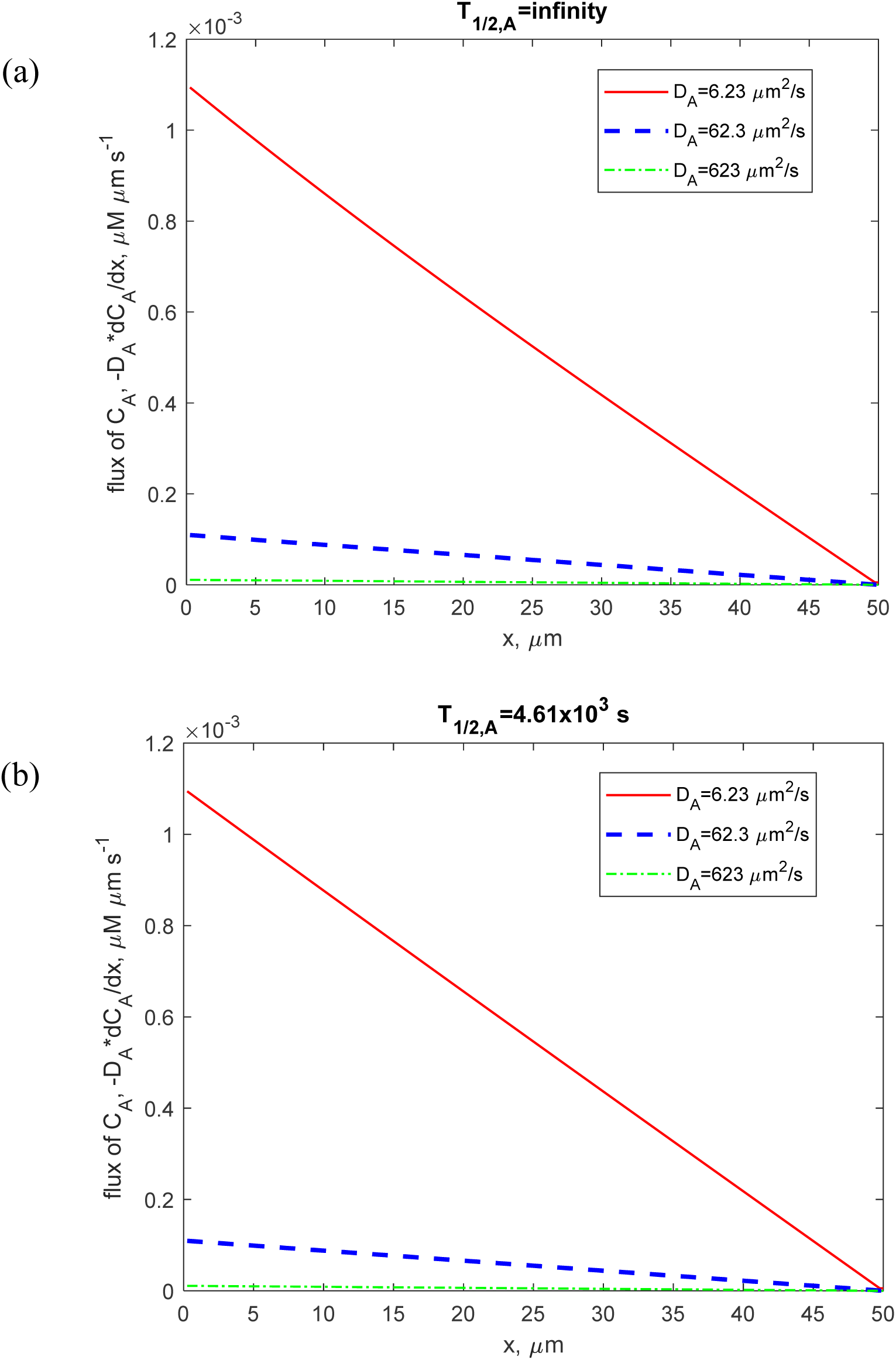
Flux of Aβ monomers as a function of the distance from the surface that releases Aβ monomers (such as the cell membrane). The two panels display the results for two different scenarios: (a) *T*_1/2,*A*_ →∞, (b) *T*_1/2,*A*_ = 4.61×10^3^ s. The computational results are presented at *t* = 2.28×10^8^ s = 7.23 years.

**Fig. S3.**
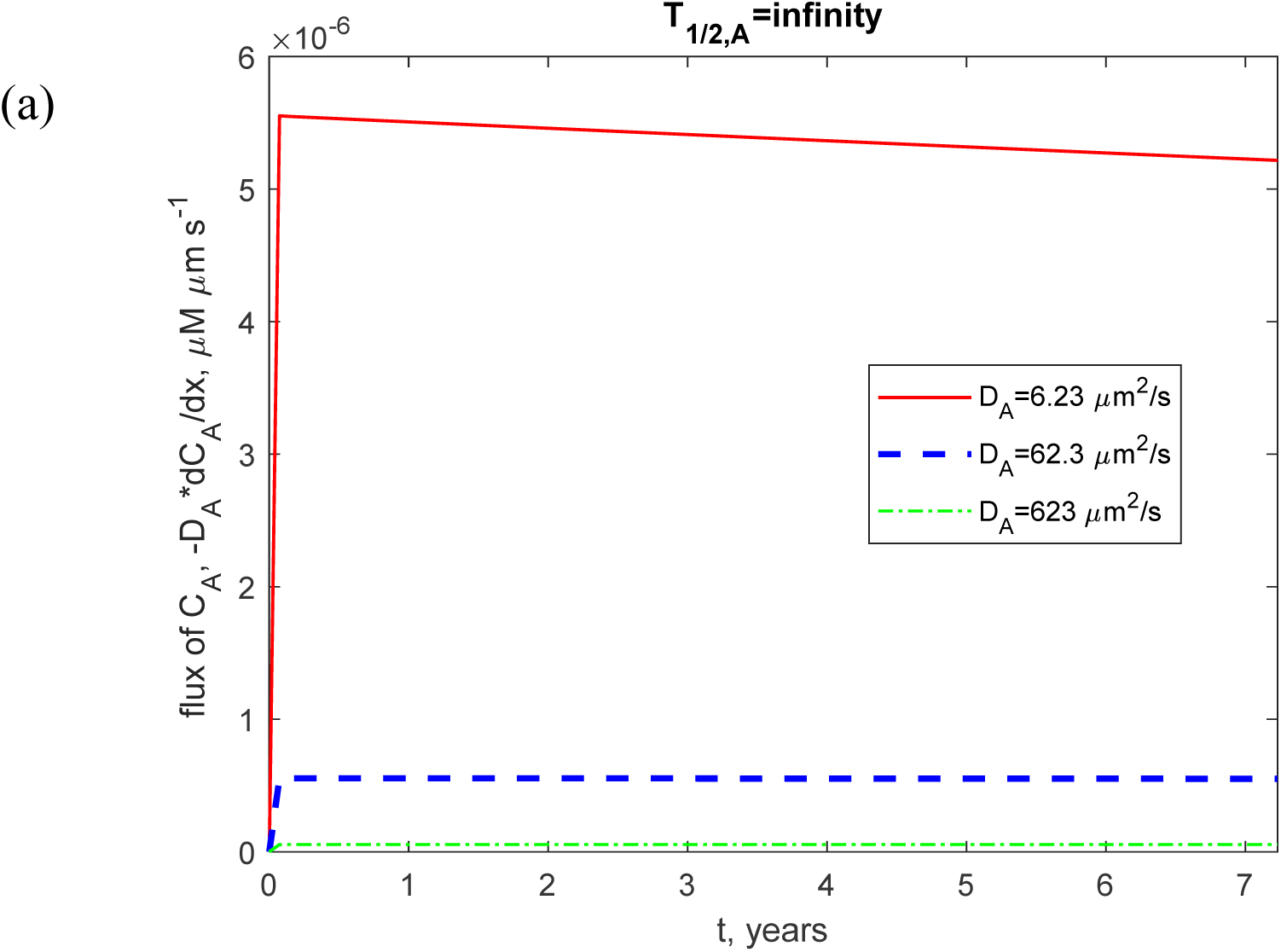
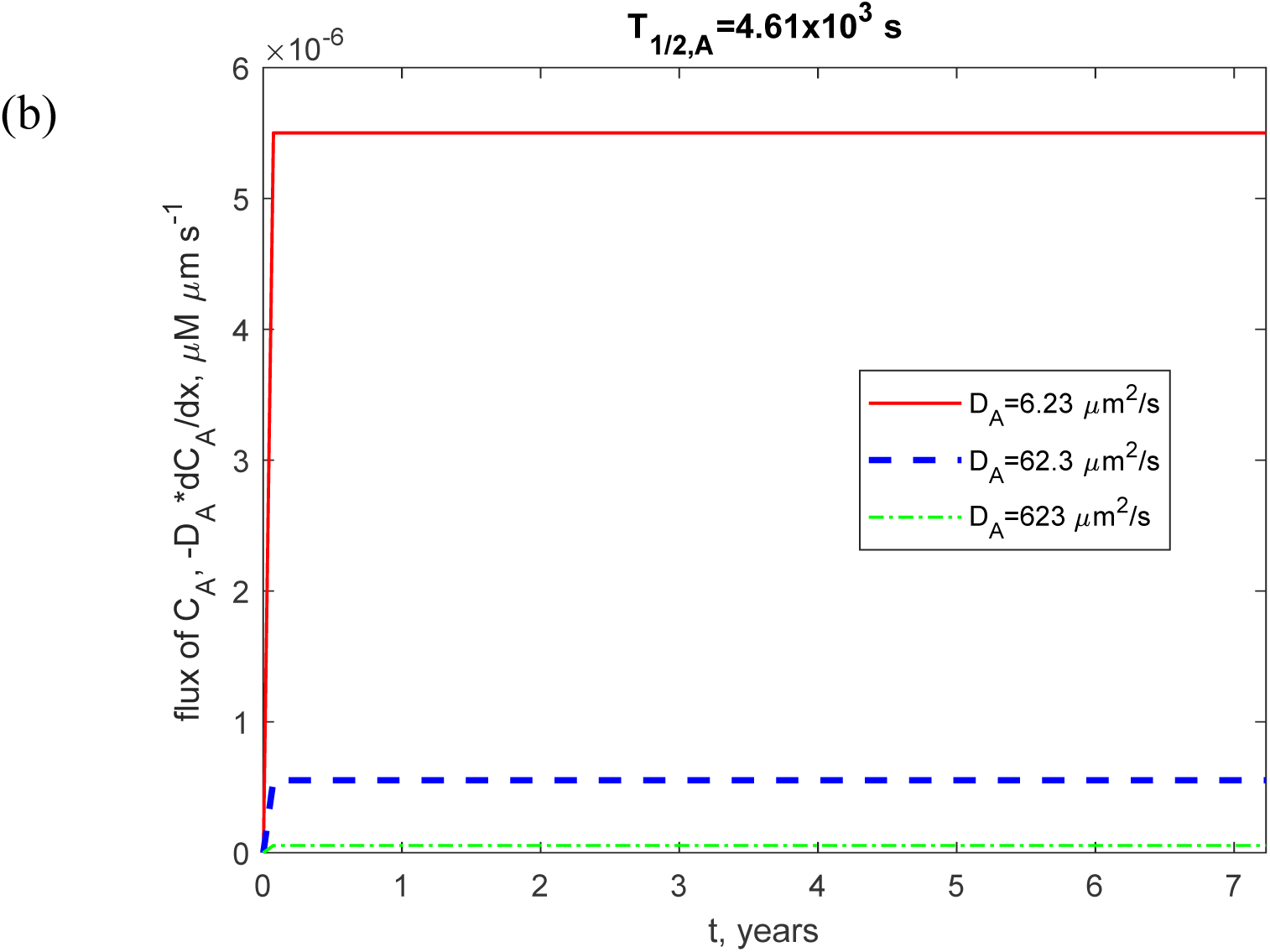
Flux of Aβ monomers as a function of time (in years). The two panels display the results for two different scenarios: (a) *T*_1/2_, *_A_* →∞, (b) *T*_1/2_,*_A_* = 4.61×10^3^ s. The computational results are presented at the right-hand side boundary of the CV (at *x=L*).

